# An Approach to Automatically Label & Order Brain Activity/Component Maps

**DOI:** 10.1101/2020.08.31.275578

**Authors:** Mustafa S. Salman, Tor D. Wager, Eswar Damaraju, Vince D. Calhoun

## Abstract

Functional magnetic resonance imaging (fMRI) is a brain imaging technique which provides detailed insights into brain function and its disruption in various brain disorders. fMRI data can be analyzed using data-driven or region-of-interest based methods. The data-driven analysis of brain activity maps involves several steps, the first of which is identifying whether the maps capture what might be interpreted as intrinsic connectivity networks (ICNs) or artifacts. This is followed by linking the ICNs to known anatomical and/or functional parcellations. Optionally, as in the study of functional network connectivity (FNC), rearranging the connectivity graph is also necessary for systematic interpretation. Here we present a toolbox that automates all these processes under minimal or no supervision with high accuracy. We provide a pretrained cross-validated elastic-net regularized general linear model for the noisecloud toolbox to separate the ICNs from artifacts. We include several well-known anatomical and functional parcellations from which researchers can choose to label the activity maps. Finally, we integrate a method for maximizing the within-domain modularity to generate a more systematically structured FNC matrix. We improve upon and integrate existing techniques and new methods to design this toolbox which can take care of all the above needs. Specifically, we show that our pretrained model achieves 89% accuracy and 100% precision at classifying ICNs from artifacts in a validation dataset. Researchers are generating brain imaging data and analyzing brain activity at an ever-increasing rate. The Autolabeller toolbox can help automate such analyses for faster and reproducible research.

## 1 Introduction

Worldwide brain imaging studies are generating diverse and large sample data at a rapidly increasing rate. Broad, collaborative efforts such as the Brain Research through Advancing Innovative Neurotechnologies (BRAIN) initiative are funding projects that produce massive amounts of data Insel et al. (2013); Mott et al. (2018). But the potential benefits of such efforts cannot be fully realized without tools that can easily share, pool, and analyze the data. It is also critical to develop and automate tools to process and analyze the data to advance neuroscience.

Functional magnetic resonance imaging (fMRI) is a widely-used technique for imaging human brain activity. fMRI can localize the variation in blood-oxygen-level-dependent (BOLD) response due to tasks or external stimuli, or at rest. However, the signal measured by fMRI is mixed with non-neural sources of variability, such as head motion, thermal noise in electrical circuits used for magnetic resonance imaging (MRI) signal reception, instrumental drifts, and hardware instability Caballero-Gaudes and Reynolds (2017). There are further physiological sources of noise affecting the MRI signal, such as cardiac and respiratory physiological noise, arterial carbon dioxide (*CO*_2_) concentration, blood pressure and cerebral autoregulation, and vasomotion Murphy et al. (2013). De-noising methods for fMRI data can be broadly divided into two categories-those that use external physiological recordings or estimates of events which might produce artifacts (e.g. motion) and those that use data-driven techniques. An example of the former is RETROICOR, in which low-order Fourier series are fit to the volume based on the time of acquisition corresponding to the phase of the cardiac and respiratory cycle Glover et al. (2000). But the precision of these methods depends on the availability and quality of the physiological measurements that are also not related to other common artifacts such as thermal noise and head movement. Data-driven approaches, on the other hand, make minimal to no assumptions about the relationship between the sources of noise and the resulting change in MRI signal and can be effective in mitigating multiple types of artifacts at once Caballero-Gaudes and Reynolds (2017). The next few paragraphs will examine different algorithmic processes within a data-driven analysis framework which the developed Autolabeller toolbox aims to automate.

Independent component analysis (ICA) is a widely used approach for de-noising fMRI and provides a powerful technique for decomposing fMRI data into maximally independent spatial components Mckeown et al. (1998). Although it is difficult to establish the correspondence between single-subject independent components (ICs) within a study, the group ICA approach can overcome this limitation Calhoun et al. (2001); Calhoun and Adali (2012). Complimentary to a fully data-driven method, a spatially constrained ICA approach can be used with ICNs determined a priori, which is especially useful for scaling up the analysis Du et al. (2016, 2020). In contrast, a fully data-driven ICA approach requires an additional identification/labelling process. In a typical ICA decomposition, some components clearly indicate BOLD signals, whereas the others indicate artifactual processes. Nevertheless, fMRI analyses can be meaningfully interpreted only after identifying, subtracting, or regressing out the artifacts. Manually classifying the ICs is time-consuming, hard to reproduce, and requires expertise Kelly et al. (2010). Therefore, several approaches have been developed for automatic classification using spatiotemporal features of the ICs. Automatic classifier types can include linear discriminant analysis, K-nearest neighbor, clustering methods, naive Bayes, sparse logistic regression, support vector machine (SVM), decision trees, random forests, or an ensemble of classifiers Tohka et al. (2008); Sui et al. (2009); Salimi-Khorshidi et al. (2014); Griffanti et al. (2014). Temporal features used by such classifiers may include spectral and/or autoregressive properties of the IC time courses (TCs), and correlation with different regression variables etc. Spatial features may include cluster size and distribution, entropy, smoothness, and the fraction of functional activation occurring in gray matter, brain edge, white matter, ventricular cerebrospinal fluid (CSF) and major blood vessels. The number of features used by the automatic classifiers may be very high to increase robustness Sochat et al. (2014); Salimi-Khorshidi et al. (2014) or very low to detect a certain type of artifact such as motion Pruim et al. (2015).

Correlating the focus of activation in functional imaging studies with structural/anatomical information is important for interpreting results. There are various anatomical atlases, methods, and tools available for this purpose. One of the most well-known atlases is the AAL atlas, which used a volumetric labeling technique to manually separate region of interests (ROIs) for a single subject brain in common stereotaxic space Tzourio-Mazoyer et al. (2002). Fig. 1 shows a mosaic view of the AAL atlas. Two more versions of the AAL atlas have been developed recently which provide alternative parcellations of the orbitofrontal cortex and add several brain areas previously not defined Rolls et al. (2015, 2020). Several tools exist for integrating anatomical parcellations and functional imaging studies. One such tool is SPM12 Anatomy, which is a toolbox for the Statistical Parametric Mapping (SPM12) software. This software provides several measures for establishing the degree of correspondence between anatomical regions and foci of functional activity, including different quantitative terms such as cluster labeling, the relative extent of activation, local maxima labeling, and relative signal change within microstructurally defined areas Eickhoff et al. (2005). Cluster labeling, which is calculated as the percentage of the overlapping voxels with an ROI relative to the total number of activated voxels in the cluster, is also used in the Autolabeller toolbox, with other available options being Pearson correlation and Matthews correlation.

**Figure 1:**
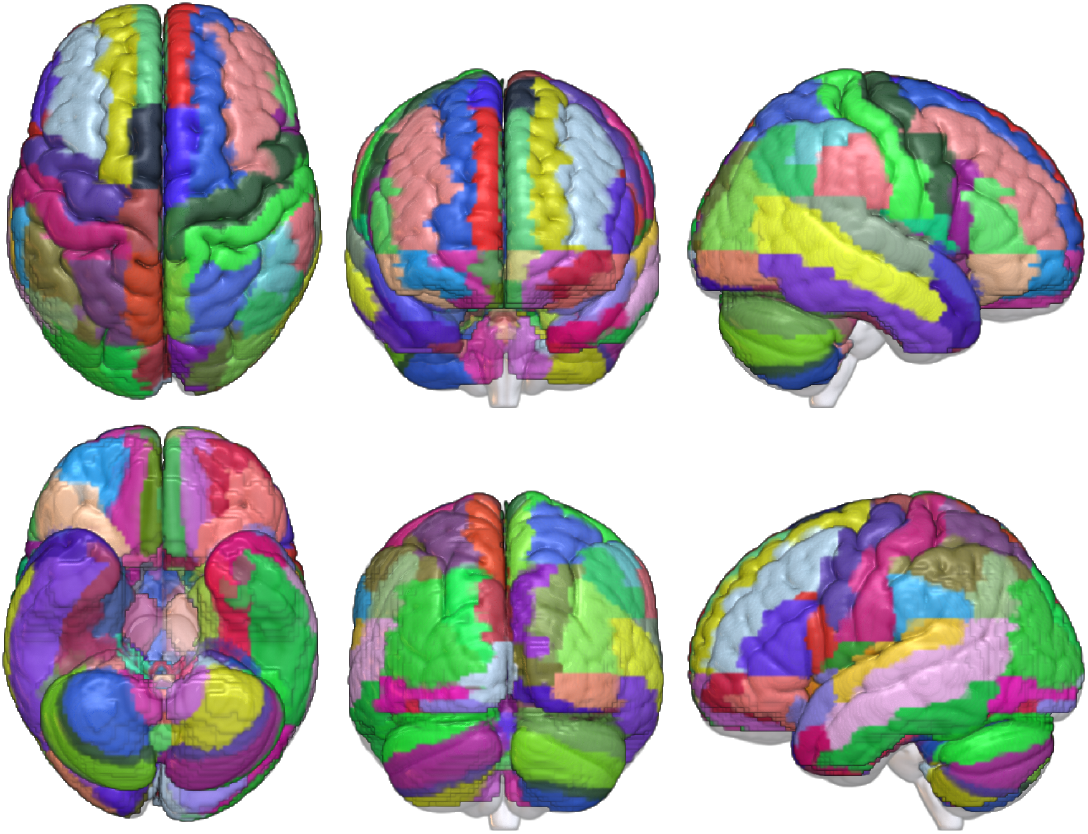
Automated Anatomical Labeling (AAL) anatomical parcellation of the brain Tzourio-Mazoyer et al. (2002); Rolls et al. (2015)

Another important aspect of fMRI studies is to examine the functional architecture of the human brain. Intrinsic connectivity networks (ICNs) derived using ICA can be divided into those involved in higher-order functions such as default mode network (DMN), central executive, and salience networks as well as externally driven sensory and motor processing such as visual and sensorimotor networks Damoiseaux et al. (2006); Smith et al. (2009); Doucet et al. (2011). Any of the several popular functional atlases can be used to determine the functional associations of IC activation maps. Yeo et al. used a clustering approach to identify functionally coupled networks across the cerebral cortex and released two well-known parcellations (7 networks and 17 networks) Yeo et al. (2011). Gordon et al. generated a parcellation of putative cortical areas using resting-state functional correlations Gordon et al. (2016). Inter-atlas variability can be concerning and Doucet et al. attempted to address this by providing the Consensual Atlas of REsting-state Networks (CAREN) atlas based on some of the most reliable atlases Doucet et al. (2019). Fig. 2 shows mosaic views of these atlases.

**Figure 2:**
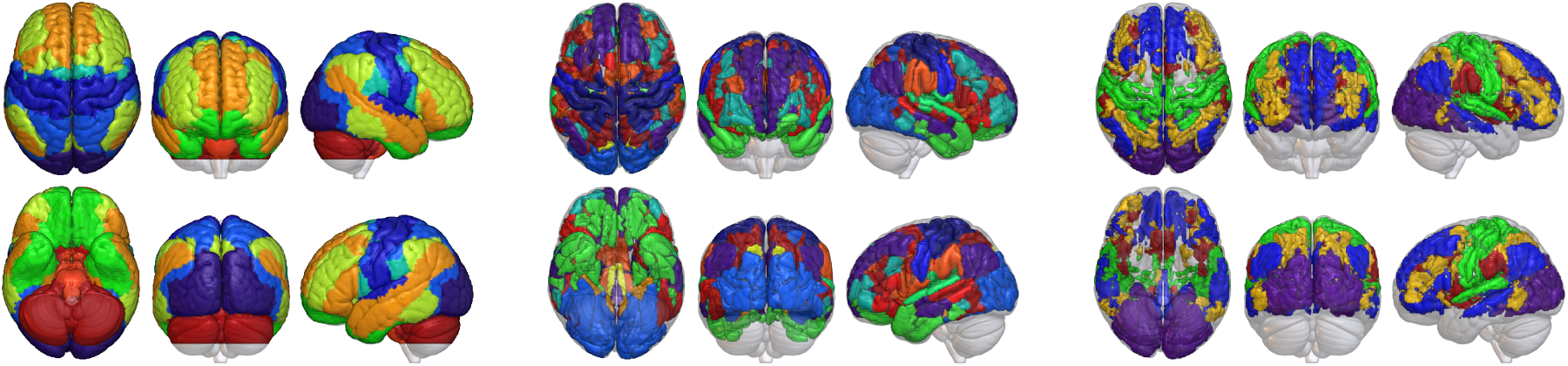
Various functional parcellations of the brain: (left) Yeo 2011 atlas with Buckner 2011 cerebellum parcellation Yeo et al. (2011); Buckner et al. (2011), (middle) Gordon 2016 atlas Gordon et al. (2016), and (right) CAREN atlas Doucet et al. (2019)

Spatial ICA analysis of fMRI data is often succeeded by the study of the temporal relationship between the ICNs or functional network connectivity (FNC) Jafri et al. (2008); Calhoun and de Lacy (2017). The FNC matrix essentially constitutes the adjacency matrix of a complete, weighted, undirected graph that can be further analyzed using graph algorithms van den Heuvel and Hulshoff Pol (2010). Given a set of ICNs with different functional labels, it is desirable to reorder the graph adjacency matrix to group ICNs with the same label together. Also, it is desirable to maximize edges with higher weights closer to the main diagonal of the adjacency matrix. This allows us to easily distinguish patterns from the FNC matrix and make meaningful observations. The Brain Connectivity Toolbox (BCT) includes an algorithm for reordering FNC matrices for this purpose Rubinov and Sporns (2010).

We describe in the paper the results of our goal to design a toolbox which can automate the above tasks. As discussed already, there are existing tools that can perform each of these tasks individually. Compared to those, the Autolabeller toolbox provides three distinct advantages. First, it incorporates all of the above tasks and provides seamless integration with the widely used group ICA for fMRI toolbox (GIFT) Calhoun (2004). Second, it includes an efficient pretrained model for separating spatial ICNs from artifacts in the fMRI data. The model parameters are supplied with the toolbox and new testing data can be classified without having to re-train the model. And finally, it can also function as a standalone software (i.e., without the GIFT toolbox); the user only needs to provide fMRI volumes in an acceptable format (e.g. Neuroimaging Informatics Technology Initiative (NIfTI)) with an optional mask.

## 2 Methods

### 2.1 Data & Preprocessing

We demonstrate the functionality of the Autolabeller toolbox using three resting-state fMRI datasets that are briefly described next. We obtained the first/primary dataset from the Function Biomedical Informatics Research Network (FBIRN) phase-III study Keator et al. (2016). This dataset includes resting-state fMRI data collected from 186 controls and 176 age and gender-matched schizophrenia patients (SZs). 162 volumes of echo planar imaging (EPI) BOLD fMRI data were collected on 3T scanners from each subject in the eyes-closed condition. The imaging parameters were the following: FOV = 220*mm×* 220*mm* (64 *×* 64 matrix), TR = 2*s*, TE = 30*ms*, flip angle = 77^*°*^, 32 sequential ascending slices with a thickness of 4*mm* and 1*mm* skip. The data were preprocessed using the SPM12 and Analysis of Functional NeuroImages (AFNI) toolboxes Cox (1996); Friston (2007). The initial 6 volumes from each scan were discarded to allow for T1-related signal saturation. The signal-fluctuation-to-noise ratio (SFNR) of all subjects was calculated and rigid-body motion correction was performed using the INRIAlign toolbox in SPM12 to obtain a measure of maximum root mean square (RMS) translation Freire et al. (2002); Friedman et al. (2006). All subjects with SFNR*<* 150 and RMS translation*>* 4*mm* were excluded. The following steps were then performed as part of preprocessing: slice-timing correction to account for timing difference in slice acquisition using middle slice as the reference, despiking using AFNI 3dDespike algorithm to mitigate the effect of outliers, spatial normalization to the Montreal Neurological Institute (MNI) space, resampling to 3*mm×* 3*mm×* 3*mm* voxels, and smoothing to 6*mm* full width at half maximum (FWHM). We retained data from 314 subjects (163 controls, mean age 36.9 years, 46 females and 151 SZs, mean age 37.8 years, 37 females) for further analysis after preprocessing and quality control. Further details about the data acquisition, preprocessing and quality control can be found in prior studies Damaraju et al. (2014); Keator et al. (2016).

We used resting-state fMRI data from the Centers of Biomedical Research Excellence (COBRE) study as the second/validation dataset under the experiment Aine et al. (2017). This study contained resting-state fMRI data from 100 controls and 87 SZs. 149 volumes of T2*-weighted functional images were acquired using a gradient-echo EPI sequence with the following parameters: TR = 2*s*, TE = 29*ms*, flip angle = 75^*°*^, 33 axial slices in sequential ascending order, slice thickness = 3.5*mm*, slice gap = 1.05*mm*, field of view = 240*mm*, matrix size = 64*×* 64 and voxel size = 3.75*mm×* 3.75*mm×* 4.55*mm*. The same quality control and preprocessing procedure were applied to both primary (FBIRN) and validation (COBRE) datasets. A total of 164 subjects (82 controls, mean age 37.7 years, 19 females and 82 SZs, mean age 38 years, 17 females) were retained for further analysis.

The third and fourth validation datasets were obtained from multiple fMRI studies on large-sample controls in the Human Connectome Project (HCP) (http://www.humanconnectomeproject.org/) and the Genomic Super-struct Project (GSP) Buckner et al. (2014). Preprocessed data from the HCP project were downloaded and then resliced to 3*mm×* 3*mm×* 3*mm* spatial resolution using SPM12. As for the GSP dataset, the following steps were performed using SPM12 as part of preprocessing: rigid body motion correction, slice timing correction, warping to the standard MNI space using an EPI template, resampling to 3*mm×* 3*mm×* 3*mm* isotropic voxels, and smoothing using a Gaussian kernel with FWHM=6*mm*.

### 2.2 Deriving Spatial Maps Using Group ICA

The simplest way to use the Autolabeller is to execute it with the group ICA session information file from the GIFT software as the input Calhoun (2004). Fig. 3 outlines the basic functionality of the Autolabeller. The GIFT toolbox was used to identify spatial ICs of brain activity from four different datasets (FBIRN and COBRE, HCP and GSP) separately in prior studies Damaraju et al. (2014); Salman et al. (2017); Du et al. (2020). In spatial group ICA, preprocessed fMRI data of each subject are first reduced from time *×* voxel dimension to *PC*1*×* voxel dimension using principal component analysis (PCA), where *PC*1 is the number of principal components (PCs).

**Figure 3:**
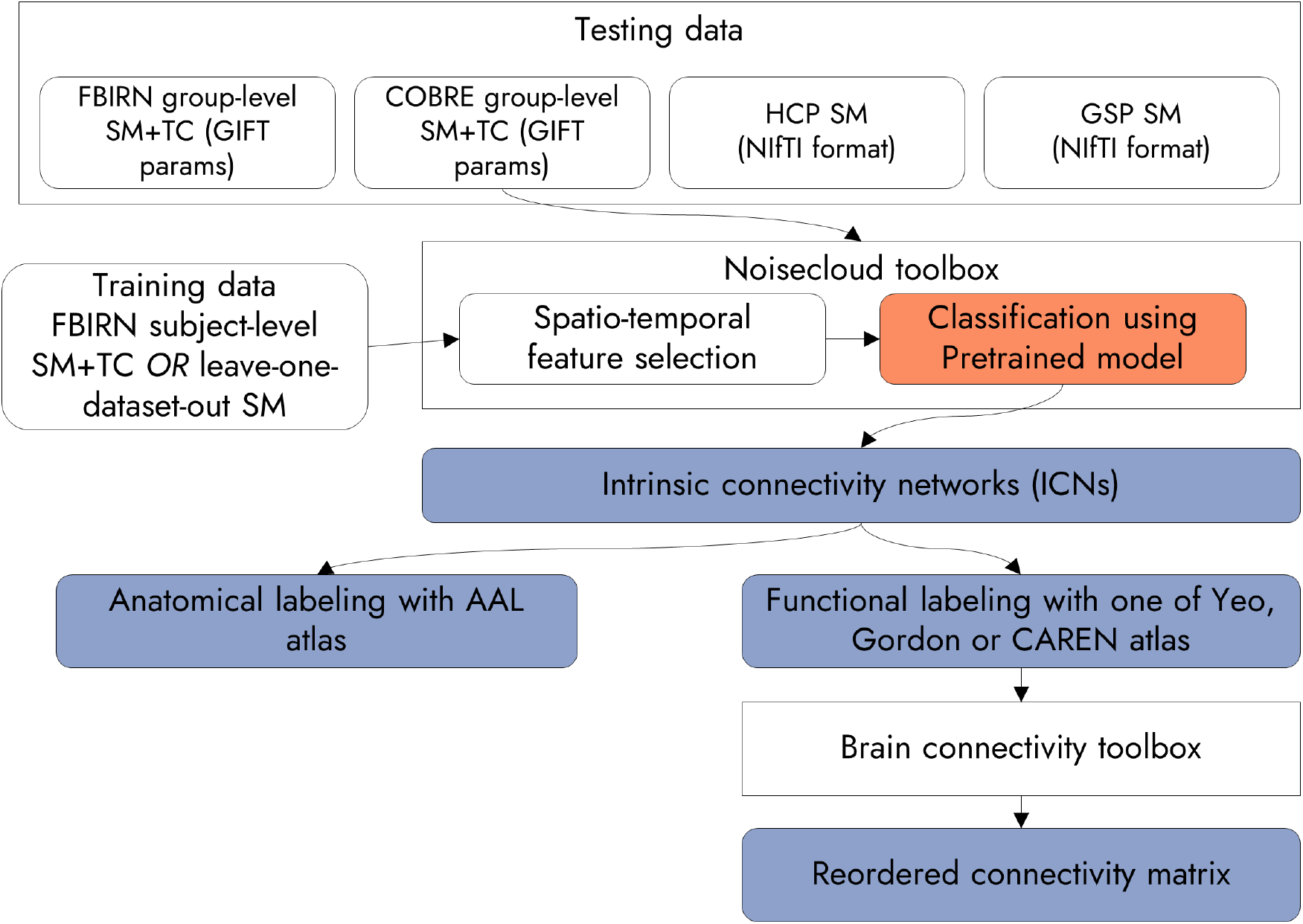
Flowchart of our analysis using the Autolabeller toolbox. SM stands for spatial map and TC for time course. Three sets of inputs are used, two of them are group ICA results from the GIFT toolbox, and one in NIfTI format. The integration of noisecloud and BCT toolboxes is shown Sochat et al. (2014); Rubinov and Sporns (2010). A pretrained model for the noisecloud toolbox, a contribution of this work, is indicated in orange. The output from the Autolabeller toolbox is indicated in purple.

Then all subjects’ data are concatenated along the time or *PC*1 dimension, and the data are reduced again to *PC*2 *×* voxel dimension using PCA. Subsequently, spatial ICs are identified from the reduced data using an ICA algorithm, such as infomax Bell and Sejnowski (1995). It is also necessary to perform the group ICA post-processing step in the GIFT toolbox by generating a report, as it also computes the FNC matrix. The aggregate ICN spatial maps (SMs) and corresponding ground truth labels of all four studies under consideration are included with the Autolabeller code Salman (2020).

### 2.3 Identifying ICNs

We integrated the noisecloud toolbox into the Autolabeller to distinguish ICNs from artifacts Sochat et al. (2014). The noisecloud toolbox implements an elastic-net regularized general linear model (GLM) with 10-fold cross-validation. It has to be trained with high-quality data for reasonable performance which we generated as follows. We obtained the labeled (ICN/noise) group-level IC SMs using the primary (FBIRN) dataset from prior work Damaraju et al. (2014). Next, we obtained the subject-level IC SMs from its group ICA result. The number of group-level ICs was 100, out of which 47 were ICNs, and the number of subjects was 314. Hence we had a dataset of 31400 ICs at our disposal, out of which 14758 were labeled ICNs and 16642 were artefactual components or noises.

It was computationally expensive and time-consuming to train the model using all of the data, and the performance gain was marginal. Therefore we randomly sampled 3000 volumes out of 31400, which we used as training data for the noisecloud toolbox. Approximately 47% of the training data were ICNs and the rest were noises. The TCs corresponding to the subject-level IC SMs were also fed into the model for training. If the input is a NIfTI volume instead of a GIFT parameter file, the TC features are not used and a model trained solely on SM features is used. The noisecloud toolbox extracts approximately 246 spatio-temporal features from the training data to train the model and uses corresponding features from the testing data to predict the ICN/noise label of the ICs. We used the mean group-level SMs, 100 each, of both the primary (FBIRN) and validation (COBRE) datasets as the testing data. The latest version of the noisecloud toolbox comes bundled with the GIFT toolbox, but can also be used in a standalone manner.

In addition to the above model trained with subject-level IC features, we trained another model using group-level IC features (aggregate SMs from the GIFT toolbox) from four datasets (FBIRN, COBRE, HCP, and GSP). In this case, the model was trained in a leave-one-dataset-out manner, i.e., a model was trained using group-level IC characteristics from three datasets (with 10-fold cross-validation), and then tested using the group-level IC features of the remaining dataset. The aggregate SMs have arbitrary scale which were thresholded by a value of 5.

### 2.4 Anatomical Labeling of SMs

We determined the anatomical label of a region of activation by correlating its SM with the masks of the AAL atlas regions Tzourio-Mazoyer et al. (2002); Rolls et al. (2015). At first, the testing data (FBIRN and COBRE mean group ICA components) were resampled to the same space as the AAL atlas using the SPM12 toolbox Friston (2007). Each region in the AAL atlas was converted to a binary mask. The pairwise correspondence between the AAL masks and volumes in the testing data was established using the Pearson correlation function in the Matlab software. Autolabeller provides two more options for determining the correspondence, i.e., Matthews correlation and cluster labeling. The top three (a number which the user can control) AAL region masks with the highest degree of correspondence with each volume are retained as the result.

### 2.5 Functional Labeling of SMs

We determined the functional label of a region of activation in the same manner as in anatomical labeling. The functional parcellations currently available in the Autolabeller include the Yeo 2011 functional parcellations (17 networks version) in conjunction with the Buckner functional cerebellar parcellation Yeo et al. (2011); Buckner et al. (2011), Gordon 2016 Gordon et al. (2016), and CAREN Doucet et al. (2019).

### 2.6 Reordering of the FNC Matrix

The Autolabeller integrates functionality from BCT for reordering the FNC matrix Rubinov and Sporns (2010). Specifically, we used the `reorder_mod` function from this toolbox to reorder the FNC matrix. This function can be applied to both binary and weighted as well as on directed and undirected graphs. It utilizes the graph community structure to reorder nodes so that the edges with higher weights are closer to the main diagonal of the graph adjacency matrix. Once the optimal order is determined, the output of the previous steps (artifact detection, anatomical, and functional labeling) are all automatically rearranged in the same order.

## 3 Results

### 3.1 Identifying ICNs

The Autolabeller incorporates a pretrained cross-validated elastic-net regularized GLM for separating ICNs from artifacts. Tab. 1 demonstrates the performance of the pretrained models applied to different datasets using two different training samples. The accuracy, precision and recall values for recognizing ICNs are all in percentages. The model predicted ICN labels with 85% accuracy in the primary (FBIRN) dataset and 89% accuracy in a validation (COBRE) dataset using both the SMs and TCs as features. A 100% precision in the validation dataset indicates that none of the true ICNs were wrongly labeled as artifacts. ICN labels were predicted with 89% accuracy in both the primary and the validation (COBRE) datasets using only the SMs as features. The asterisk (*) indicates that the testing data was biased, i.e., FBIRN mean or aggregate ICA components were tested using a model trained with FBIRN subject-level IC features. In the validation datasets where only the SM features were used, i.e., HCP and GSP, the accuracy was still reasonable (86% and 77% respectively).

**Table 1:** Performance in detecting ICN vs. noise using a pretrained model.

| Testing data | Training data | Training N | Train. acc. | Test. acc. | Precision | Recall |
| --- | --- | --- | --- | --- | --- | --- |
| FBIRN mean SM+TC | FBIRN sub. SM+TC | 3000 | 87.19 | 85 | 80.85 | 86.36 |
| COBRE mean SM+TC |  | 3000 | 87.19 | 89 | 100 | 76.59 |
| FBIRN agg. SM | Leave-one-dataset<br>-out SM | 300 | 74 | 89 | 80.85 | 95 |
| COBRE agg. SM |  | 300 | 71.67 | 89 | 94.44 | 79.07 |
| HCP agg. SM |  | 300 | 74.33 | 86 | 78.43 | 93.02 |
| GSP agg. SM |  | 300 | 85 | 77 | 66.67 | 85 |

### 3.2 Anatomical & Functional Labeling

Tab. 2 and 3 show sample anatomical and functional labels respectively in a tabular text format as generated by the Autolabeller. For each fMRI volume, the output indicates whether it is an ICN or not, and lists the top three highest correlated ROIs in the chosen anatomical/functional parcellation. Also, supplementary Tab. 1 & 2 list the anatomical and functional labels of the ICN SMs from the FBIRN and COBRE datasets respectively. Supplementary fig. 1 & 2 depicts the ICN SMs from the FBIRN and COBRE datasets respectively, grouped into functional domains. The tables list the labels determined by the Autolabeller as well as those from prior works (ground truths). For the FBIRN dataset, 32 out of 47 (68.08%) of both the anatomical and functional labels agree with the labeling from the prior work Damaraju et al. (2014). For the COBRE dataset, 25 out of 36 (69.44%) of both the anatomical and functional labels agree with the labeling from the prior work Salman et al. (2017).

**Table 2:**
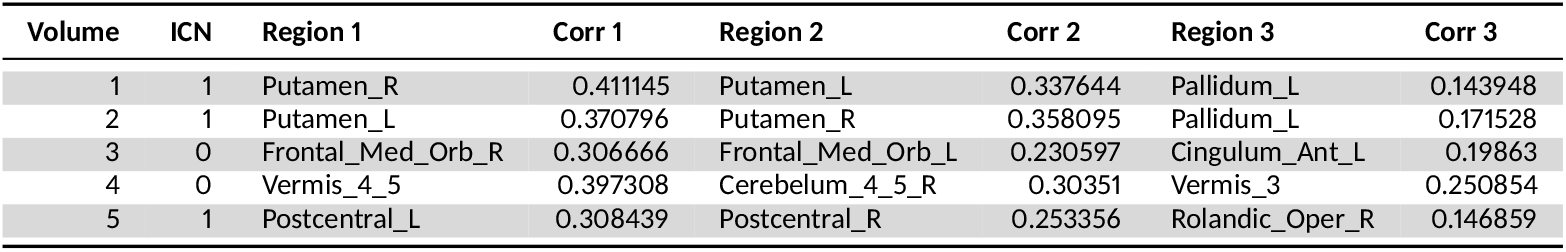
Sample FBIRN anatomical labeling output based on the AAL atlas.

**Table 3:** Sample FBIRN functional labeling output based on the Yeo 2011 atlas.

| Volume | ICN | Region 1 | Corr 1 | Region 2 | Corr 2 | Region 3 | Corr 3 |
| --- | --- | --- | --- | --- | --- | --- | --- |
| 1 | 1 | basal ganglia | 0.429328 | basal ganglia | 0.387354 | basal ganglia | 0.151536 |
| 2 | 1 | basal ganglia | 0.269627 | basal ganglia | 0.261333 | basal ganglia | 0.237799 |
| 3 | 0 | Default | 0.413814 | Limbic | 0.155148 | Default | 0.073393 |
| 4 | 0 | cerebellum | 0.509603 | cerebellum | 0.261651 | cerebellum | 0.187854 |
| 5 | 1 | Somatomotor | 0.374891 | Somatomotor | 0.373648 | cerebellum | 0.046119 |

### 3.3 Reordering of the FNC Matrix

Fig. 4 displays the FNC matrices for the unordered components (left), separate ICN and noise components (middle), and ordered ICNs (right) in the FBIRN (top) and COBRE (bottom) datasets. The ICNs on the right are grouped into functional domains based on the Yeo 2011 atlas and Buckner cerebellar parcellation. The FNC matrices are ordered by the Autolabeller to increase the modularity of the community structures within each functional domain. As such, the reordered matrices are much easier to visualize and interpret.

**Figure 4:**
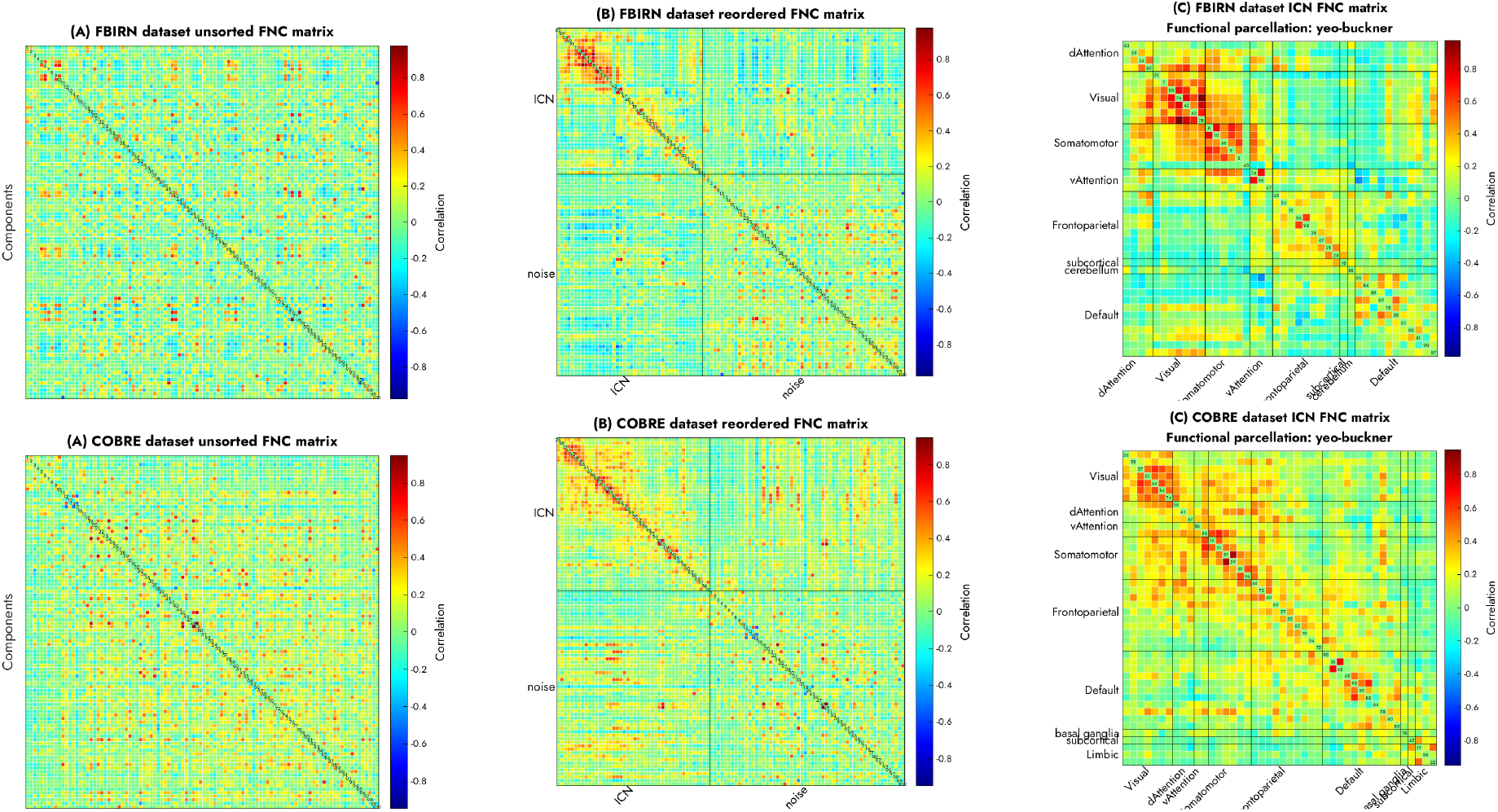
Unordered vs. automatically reordered FNC matrices. The Autolabeller separates resting-state ICNs from noise and groups the ICNs into functional domains with high modularity.

## 4 Discussion

Separating artifacts from ICNs, labeling the foci of functional activation, and modularizing the functional connectivity are some of the tasks still many researchers perform manually. Here we design a toolbox that can enable automation of these tasks for improving brain imaging studies. It provides the user with relevant anatomical and functional correlates of functional activation, which facilitate further analysis of neuroimaging results. Other advantages of the Autolabeller toolbox include the fact that it is modular in design and the user can choose which steps to run. The intermediate results are written out in a comma-delimited tabular text format that is easy to inspect, edit, and be fed back into the Autolabeller or read into any other software. The tabular text output even enables the user to use the Autolabeller output as a guideline and customize the results based on their subjective opinion. The toolbox can be seamlessly integrated with group ICA analysis in the GIFT toolbox, but can also be used in a standalone fashion with fMRI data in an acceptable format (e.g. NIfTI) and can incorporate an optional brain mask.

Next, we discuss some of the subtle differences between the Autolabeller output and the ground truths that are based on prior studies. The Top panel of Fig. 5 displays the contrast between the ICN/noise labels of some of the SMs generated by the Autolabeller. Volume 13 (top left panel) is labeled as a noisy component, but in the prior work, it is labeled as an ICN Damaraju et al. (2014). As the noisecloud toolbox uses approximately 246 spatio-temporal features, it is difficult to pinpoint what causes the model to flag this as a noisy component without a thorough feature importance analysis. But a cursory look at the noisecloud testing features file (provided with the code) indicates some distinct temporal features common in noisy components, such as low average distance between max/mean peaks and a high number of local maxima/minima in the TC. In the case of volume 67 (top right panel), it is the opposite; it is labeled as an ICN by the Autolabeller but the prior work ignores it as a noisy component. According to the Autolabeller output, it has all the opposite temporal characteristics compared to volume 13. Supplementary tables list all the volumes in FBIRN and COBRE datasets respectively which are marked as ICNs either by the Autolabeller or in prior work Damaraju et al. (2014); Salman et al. (2017). The bottom panel of Fig. 5 displays the contrast between the anatomical/functional labels of some of the SMs. Volume 7 (bottom left panel) is labeled as a bilateral calcarine component, as it has a correlation value of 0.44 with the left Calcarine ROI and 0.39 with the right Calcarine ROI. The Autolabeller output (provided with the code) lists right Lingual gyrus the 3rd highest correlated (0.13) ROI. However, in the prior work, it is labeled as right Cuneus Damaraju et al. (2014). On the other hand, volume 12 (bottom right panel) is labeled as bilateral Calcarine by both the Autolabeller and the prior work, but in terms of function, it is associated with DMN and visual networks respectively. According to the Autolabeller output, this volume is highly correlated with multiple DMN ROI of the Yeo 2011 atlas (top 3 correlation values of 0.41, 0.4, and 0.25).

**Figure 5:**
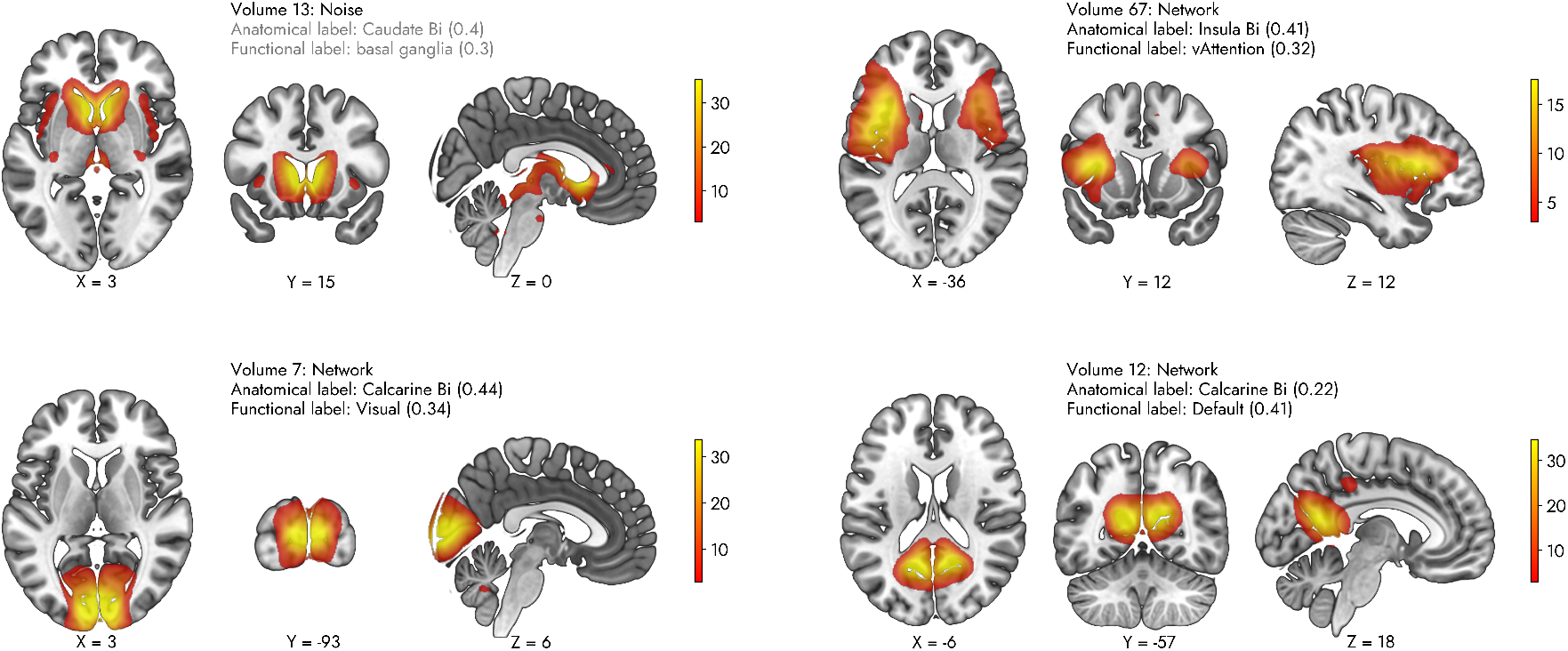
Examples of Autolabeller prediction in FBIRN dataset

We retained the noisecloud toolbox in the Autolabeller but generated a model ourselves for using in conjunction with it. The noisecloud toolbox, available as part of the GIFT toolbox, already includes an algorithm that can extract 246 different spatial and temporal features from the input data. This is one of the largest feature sets in the literature for this particular problem, which makes for a robust algorithm capable of generalizing well and detecting multiple types of noise components. Our model was trained using a subset of the subject-level SMs and TC features from the primary dataset (FBIRN). During prediction, it achieved higher accuracy in the validation dataset (COBRE) compared to the primary dataset, which proves that the model generalizes very well. Although we provide a pretrained model, the user still has the option to provide their training data, which allows for more flexibility and better sensitivity especially when a large volume of training data is available.

There are also a few limitations inherent to noisecloud integration. For instance, the noisecloud toolbox prediction is very fast with a pretrained model. However, training with new data is somewhat slow, i.e., it can take about 20 seconds to extract features from one fMRI volume by a single process on a computer. This process scales linearly and can take quite long for a large amount of training data. This issue can be mitigated by refactoring the noisecloud toolbox to leverage parallel processing. Furthermore, in our analysis, we did not regress any nuisance covariates, such as motion parameters, from the training or testing data. The noisecloud toolbox provides an option to specify such information, which should make the prediction more robust and accurate. The classification can also be repeated to determine the optimal amount of training data and further validate the classification result.

The Autolabeller integrates well-known atlases such as AAL, Yeo 2011, Gordon 2016, and CAREN for analyzing the anatomical and functional correlates of brain activity. By using these labeling, researchers can publish reproducible results instantly comparable to other studies using any of these same atlases. The results show that compared to prior work using the same primary and validation datasets, the accuracy of anatomical and functional labeling is rather low (below 70%). This is however expected because prior works used subjective criteria for labeling instead of quantitatively determining association with a known atlas Allen et al. (2014). Another limitation of the Autolabeller is a limited number of atlas choices. In particular, not every one of these may cover the entire cortical, subcortical, and cerebellar regions of the brain. The Yeo 2011 functional atlas choice in the autolabeller in fact integrates both the Yeo 2011 cortical parcellation and Buckner 2011 cerebellum parcellation. However, Gordon 2016 or CAREN atlases do not include cerebellum parcellation. We will add support for more atlases, both anatomical and functional in the future to alleviate these problems.

## 5 Code & Data Availability

The Autolabeller toolbox is available on GitHub along with the training/testing data used in this work and the supplementary tables Salman (2020). It can be used in conjunction with the GIFT toolbox, available at https://trendscenter.org/software/.

