## Supplementary data for "An Approach to Automatically Label & Order Brain Activity/Component Maps"

### 1 Supplementary Materials

#### 1.1 Intrinsic connectivity networks (ICNs)

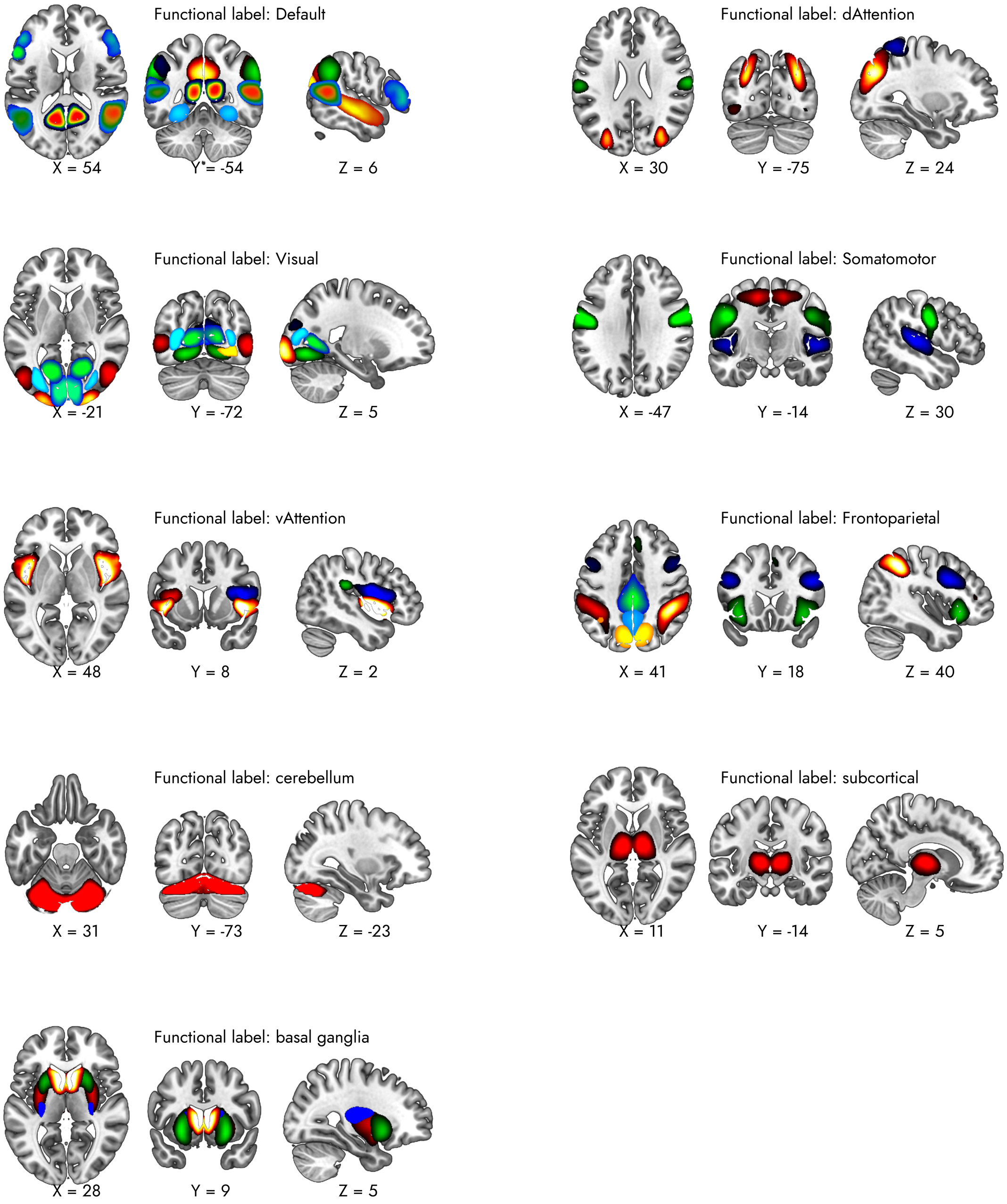


Figure 1: ICNs estimated from the Function Biomedical Informatics Research Network (FBIRN) dataset, grouped into functional domains based on the Yeo 2011 atlas

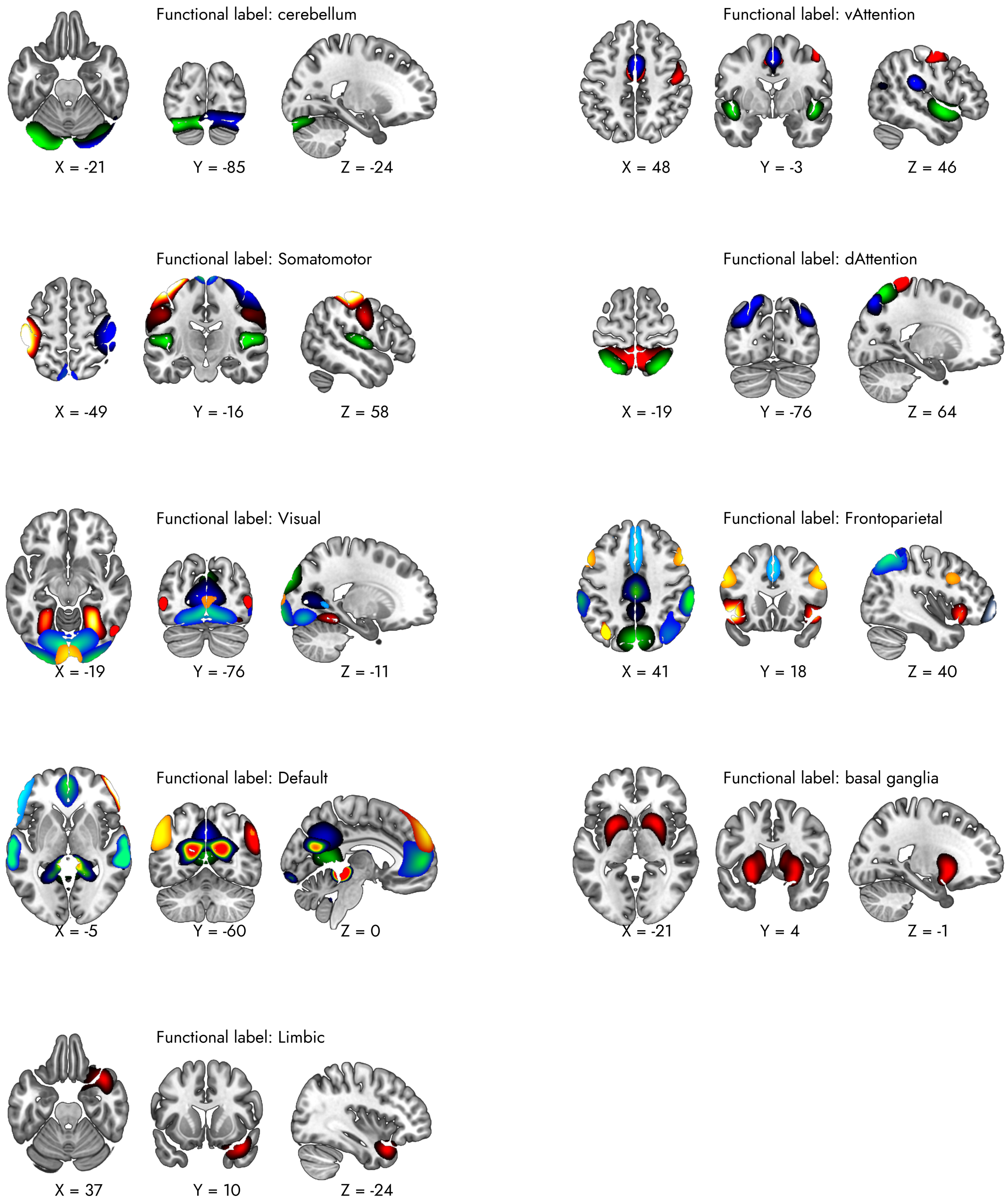


Figure 2: ICNs estimated from the Centers of Biomedical Research Excellence (COBRE) dataset, grouped into functional domains based on the Yeo 2011 atlas

#### 1.2 Anatomical & Functional Labels

Table 1: FBIRN Dataset Anatomical & Functional Labels. Only the volumes labeled as networks either by the Autolabeller or Damaraju et al. (2014) are listed. Volumes labeled as artifacts by either are marked as N/A.

| | Anatomical labels | | Functional labels | |
|---|---|---|---|---|
| **Volume** | **Autolabeller** | **Damaraju et al. (2014)** | **Autolabeller'** | **Damaraju et al. (2014)'** |
| **1** | Putamen Bi | Putamen | basal ganglia | Subcortical |
| **2** | Putamen Bi | Putamen | basal ganglia | Subcortical |
| **5** | Postcentral Bi | Ventral Motor | Somatomotor | Somatomotor |
| **6** | Postcentral R | Motor R | Somatomotor | Somatomotor |
| **7** | Calcarine Bi | Ventral Cuneus | Visual | Visual |
| **9** | Paracentral Biobule Bi | Paracentral | Somatomotor | Somatomotor |
| **10** | Postcentral L | Motor L | Somatomotor | Somatomotor |
| **12** | Calcarine Bi | Calcarine | Default | Visual |
| **13** | N/A | Caudate | basal ganglia | Subcortical |
| **18** | Thalamus Bi | Thalamus | subcortical | Subcortical |
| **20** | Occipital Inf Bi | Occipital Inf | Visual | Visual |
| **21** | Frontal Mid Bi | Frontal Mid | Frontoparietal | Frontoparietal |
| **24** | Parietal Sup Bi | Parietal Sup Bi | dAttention | Attention |
| **28** | Frontal Inf Orb Bi | Anterior Insula | Frontoparietal | Frontoparietal |
| **30** | Precuneus Bi | Posterior Cingulate | Default | Default Mode |
| **34** | Frontal Inf Oper Bi | Precentral | Frontoparietal | Frontoparietal |
| **35** | Cuneus Bi | Precuneus | Frontoparietal | Attention |
| **40** | Precuneus Bi | Precuneus | Frontoparietal | Attention |
| **41** | Insula Bi | Posterior Insula | vAttention | Frontoparietal |
| **42** | Occipital Mid Bi | Temporal Mid/Occipital Mid | Visual | Visual |
| **43** | Calcarine Bi | Calcarine | Visual | Visual |
| **46** | Cerebelum 6 Bi | Cerebelum | cerebellum | Cerebellum |
| **47** | Cingulum Mid Bi | Posterior Cingulate | Frontoparietal | Default Mode |
| **51** | Temporal Mid Bi | Temporal Sup | Default | Auditory |
| **53** | Cingulum Ant Bi | Anterior Cingulate | Default | Default Mode |
| **57** | Fusiform Bi | Fusiform | Default | Visual |
| **58** | Temporal Sup Bi | Temporal Sup | Somatomotor | Auditory |
| **59** | SupraMarginal Bi | Bi Postcentral | dAttention | Somatomotor |
| **60** | Occipital Mid Bi | Occipital Sup | dAttention | Visual |
| **61** | Temporal Mid L | Temporal Mid L/Frontal Inf | Default | Frontoparietal |
| **63** | Fusiform Bi | Temporal Inf | dAttention | Attention |
| **64** | N/A | N/A | dAttention | N/A |
| **65** | Frontal Inf Tri Bi | Frontal Inf | Default | Frontoparietal |
| **66** | Parietal Inf R | Parietal Inf | Frontoparietal | Attention |
| **67** | Insula Bi | N/A | vAttention | N/A |
| **69** | Frontal Sup Medial Bi | dMPFC | Default | Frontoparietal |
| **73** | Temporal Inf Bi | N/A | Default | N/A |
| **74** | Supp Motor Area Bi | Supp Motor Area | vAttention | Somatomotor |
| **75** | N/A | Substantia Nigra | subcortical | Subcortical |
| **76** | Lingual Bi | Lingual | Visual | Visual |
| **78** | Cuneus Bi | Cuneus | Visual | Visual |
| **80** | Temporal Mid Bi | Temporal Mid | Default | Attention |
| **84** | Angular L | Angular L | Default | Default Mode |
| **88** | Cerebelum Crus1 Bi | Cerebelum | cerebellum | Cerebellum |
| **89** | Angular Bi | Parietal Inf | Default | Attention |
| **90** | Occipital Mid Bi | Angular | Default | Default Mode |
| **91** | Occipital Inf R | Occipital Mid R | Visual | Visual |
| **94** | Parietal Inf L | Parietal Inf L | Frontoparietal | Attention |
| **95** | Angular L | Angular L | Default | Default Mode |
| **96** | SupraMarginal Bi | SupraMarginal | vAttention | Attention |

Table 2: COBRE Dataset Anatomical & Functional Labels. Only the volumes labeled as networks either by the Autolabeller or Salman et al. (2017) are listed. Volumes labeled as artifacts by either are marked as N/A.

| | **Anatomical labels** | | **Functional labels** | |
|---|---|---|---|---|
| **Volume** | **Autolabeller** | **Salman et al. (2017)** | **Autolabeller'** | **Salman et al. (2017)'** |
| **11** | Postcentral Bi | Postcentral Bi | Somatomotor | Somatomotor |
| **17** | Rectus Bi | N/A | Limbic | N/A |
| **18** | Putamen Bi | Putamen Bi | basal ganglia | Subcortical |
| **19** | Supp Motor Area Bi | N/A | vAttention | N/A |
| **21** | Postcentral R | Postcentral R | Somatomotor | Somatomotor |
| **22** | Cerebelum Crus1 R | Lingual R | cerebellum | Visual |
| **26** | Occipital Inf Bi | Occipital Bi | Visual | Visual |
| **29** | Postcentral L | Postcentral L | Somatomotor | Somatomotor |
| **30** | Paracentral Lobule Bi | N/A | Somatomotor | N/A |
| **33** | Cerebelum Crus1 Bi | Cerebelum Bi | cerebellum | Cerebellum |
| **34** | Cuneus Bi | Cuneus Bi | Visual | Visual |
| **35** | Frontal Sup Medial Bi | N/A | Default | N/A |
| **36** | Supp Motor Area Bi | N/A | dAttention | N/A |
| **37** | Lingual Bi | Calcerine Bi | Visual | Visual |
| **40** | Cuneus Bi | Precuneus Bi | Frontoparietal | Default mode |
| **41** | Parietal Sup Bi | Precuneus Bi | dAttention | Default mode |
| **42** | Olfactory Bi | N/A | subcortical | N/A |
| **46** | Precuneus Bi | Precuneus Bi | Frontoparietal | Default mode |
| **48** | Frontal Sup Bi | N/A | Frontoparietal | N/A |
| **50** | Cingulum Ant Bi | Cingulum Ant Bi | Default | Default mode |
| **52** | Frontal Inf Oper Bi | Frontal Inf Oper Bi | Frontoparietal | Cognitive control |
| **56** | Occipital Mid Bi | Occipital Bi | dAttention | Visual |
| **57** | Cingulum Mid Bi | N/A | Frontoparietal | N/A |
| **58** | Calcarine Bi | Calcarine Bi | Visual | Visual |
| **61** | Fusiform Bi | Fusiform Bi | Visual | Visual |
| **63** | Calcarine Bi | Calcarine Bi | Visual | Visual |
| **65** | Thalamus Bi | N/A | subcortical | N/A |
| **67** | SupraMarginal Bi | SupraMarginal Bi | Frontoparietal | Somatomotor |
| **69** | Vermis 4 5 | Precentral Bi | Default | Default mode |
| **70** | ParaHippocampal Bi | N/A | Default | N/A |
| **71** | Parietal Sup Bi | Parietal Sup Bi | dAttention | Somatomotor |
| **73** | Precuneus Bi | Mid Cingulate Bi | Frontoparietal | Default mode |
| **74** | Temporal Mid Bi | Temporal Mid Bi | Visual | Visual |
| **75** | Temporal Sup Bi | Heschl's Bi | Somatomotor | Auditory |
| **78** | Temporal Mid Bi | Temporal Mid Bi | Default | Auditory |
| **79** | Parietal Inf L | Parietal Sup L | Frontoparietal | Somatomotor |
| **80** | Precentral Bi | N/A | vAttention | N/A |
| **81** | Paracentral Lobule R | Paracentral lobule R | Somatomotor | Somatomotor |
| **82** | Frontal Sup Medial Bi | Frontal Sup Medial Bi | Default | Default mode |
| **83** | Parietal Inf R | Angular R | Frontoparietal | Default mode |
| **84** | Temporal Pole Sup Bi | N/A | Limbic | N/A |
| **85** | Angular L | Angular L | Default | Default mode |
| **87** | Lingual Bi | N/A | Visual | N/A |
| **88** | SupraMarginal Bi | Temporal Sup Bi | vAttention | Auditory |
| **89** | Insula Bi | Insula Bi | Frontoparietal | Cognitive control |
| **90** | Angular R | Temporal Sup R | Default | Default mode |
| **93** | Precuneus Bi | Precuneus Bi | Default | Default mode |
| **94** | Frontal Mid Orb R | Frontal Sup Medial R | Frontoparietal | Cognitive control |
| **96** | Paracentral Lobule L | Paracentral lobule L | Somatomotor | Somatomotor |
